# A scalable method for estimating the regional polygenicity of complex traits

**DOI:** 10.1101/2020.01.15.908095

**Authors:** Ruth Johnson, Kathryn S. Burch, Kangcheng Hou, Mario Paciuc, Bogdan Pasaniuc, Sriram Sankararaman

## Abstract

A key question in human genetics is understanding the proportion of SNPs modulating a particular phenotype or the proportion of susceptibility SNPs for a disease, termed *polygenicity*. Previous studies have observed that complex traits tend to be highly polygenic, opposing the previous belief that only a handful of SNPs contribute to a trait. Beyond these genome-wide estimates, the distribution of polygenicity across genomic regions as well as the genomic factors that affect regional polygenicity remain poorly understood. A reason for this gap is that methods for estimating polygenicity utilize SNP effect sizes from GWAS. However, estimating regional polygenicity from GWAS effect sizes involves untangling the correlation between SNPs due to LD, leading to intractable computations for even a small number of SNPs. In this work, we propose a scalable method, BEAVR, to estimate the regional polygenicity of a trait given marginal effect sizes from GWAS and LD information. We implement a Gibbs sampler to estimate the posterior distribution of the regional polygenicity and derive a fast, algorithmic update to circumvent the computational bottlenecks associated with LD. The runtime of our algorithm is 𝒪(*MK*) for *M* SNPs and *K* susceptibility SNPs, where the number of susceptibility SNPs is typically *K* ≪ *M*. By modeling the full LD structure, we show that BEAVR provides unbiased estimates of polygenicity compared to previous methods that only partially model LD. Finally, we show how estimates of regional polygenicity for BMI, eczema, and high cholesterol provide insight into the regional genetic architecture of each trait.

## 1 Introduction

Genome-wide association studies (GWAS) have identified tens of thousands of genetic variants in the genome that are associated with an increased risk for many diseases, where the effect of a variant on a trait is estimated by its marginal regression coefficient. Due to pervasive correlations among genotypes at single nucleotide polymorphisms (SNPs) – a phenomenon known as linkage disequilibrium (LD) – a SNP that is estimated to be significantly associated with a trait could actually have no true effect on the trait. We refer to SNPs that have a nonzero effect on a trait as *susceptibility SNPs* and the proportion of susceptibility SNPs for a trait as the *polygenicity* of the trait. Accurately estimating polygenicity can aid in understanding the genetic basis of a trait, as well as estimating the upper bound for the accuracy of phenotype prediction [1] from genetic data, the response of a trait under natural selection [2, 3], and the power to detect novel genetic associations from GWAS [4].

Genetic architecture has been analyzed both at the genome-wide and regional level, where the latter refers to analyses done within a given genomic region. Previous work has already highlighted the utility of regional analyses through the partitioning of heritabilities across regions of the genome [5, 6], where *heritability* refers to the amount of phenotypic variation explained by the genetic variation for a trait [7]. Similar to how genome-wide heritability can be partitioned into regional heritability, genome-wide polygenicity can be partitioned into *regional polygenicities*, defined here as the number of susceptibility SNPs within a given region of the genome. Investigating regional polygenicity can be means to identify regions that contribute a disproportionately large number of susceptibility SNPs and provides novel insight into the physical distribution of susceptibility SNPs across the genome.

It is not straightforward to infer the polygenicity of a trait directly from GWAS effects. Due to LD and noise from the regression performed in GWAS, all effect sizes estimated from GWAS are non-zero, but not every SNP is truly a susceptibility SNP. Furthermore, even SNPs that meet genome-wide significance (p-value < 5*e*-8) are not guaranteed to have a true effect for the trait due to LD. Estimating polygenicity also presents a computationally challenging inference problem. Because the marginal effects from GWAS are correlated, estimating polygenicity from GWAS while accounting for LD requires fully conditioning on the “susceptibility status” of every SNP. Performing inference under this model involves explicitly enumerating all possible configurations of susceptibility SNPs. This creates an exponential search space of 2^*M*^, where M is the number of SNPs, which is intractable even when analyses are within small regions in the genome. To circumvent the large computational bottleneck, existing methods that estimate polygenicity from GWAS do not explicitly condition on the susceptibility status of every SNP[4]. Thus, we expect these methods to lead to a downward bias when estimating polygenicity since only partially modeling the LD structure prevents these methods from fully re-capturing SNPs’ effects that have been spread throughout the region due to LD. At a regional level, we expect the impact of a downward bias to be more pronounced since underestimating the polygenicity in a region with a small fraction of susceptibility SNPs can be the difference between estimating the absence of susceptibility SNPs or the presence of only a few.

### 1.1 Novel contributions

In this work, we propose a statistical framework to estimate the regional polygenicities of a complex trait from GWAS while fully modeling the correlation structure due to LD. We present a fully generative Bayesian statistical model that estimates, for a given region, the regional polygenicity. Our model inherently allows for a variety of genetic architectures as it does not make prior assumptions on the number of susceptibility SNPs for a trait. We propose Bayesian EstimAtion of Variants in Regions (BEAVR) which relies on Markov Chain Monte Carlo (MCMC) to estimate the posterior distribution of the regional polygenicities of a trait. A straightforward implementation of the MCMC sampler still presents a computational bottleneck since each iteration of the sampler is 𝒪(*M* ^2^) due to the full conditioning on each SNP, where M is the number of SNPs. To address this, we introduce a new inference algorithm that leverages the intuition that the majority of SNPs are not susceptibility SNPs. This algorithm allows us to perform efficient inference that scales approximately linearly with the number of SNPs analyzed in a region. Through comprehensive simulations and an analysis of BMI, eczema, and high cholesterol from the UK Biobank [8], we show that our method can accurately estimate regional polygenicity across many settings and provides insight into the genetic architecture for a variety of traits that is consistent with previous knowledge as well as provides novel information about the physical distribution of susceptibility SNPs across the genome.

## 2 Materials and Methods

### 2.1 Generative model

A standard approach to describe a phenotype is through a linear model. We assume that the trait measured in individual *i, y*_*i*_, is a linear function of standardized genotypes ***x***_***i***_ = (*x*_*i*,1_, …, *x*_*i,M*_) measured across *M* SNPs with true SNP effect sizes ***β*** = (*β*_1_, …, *β*_*M*_)^*T*^ and independent additive noise term *ϵ*_*i*_.

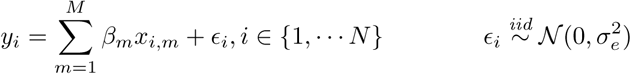

To model whether or not a SNP has a true effect for a trait, we impose a spike-and-slab prior on the true effect sizes, ***β*** [9]. To model this, we use an indicator variable to represent each SNP’s susceptibility status, where the (*M* × 1) vector ***c*** = (*c*_1_, …, *c*_*M*_)^*T*^ denotes the susceptibility status indicator vector across all *M* SNPs. Here, *c*_*m*_ = 1 if SNP *m* is a susceptibility SNP with probability *p* and 0 otherwise with probability (1 − *p*). Thus, *p* is interpreted as the proportion of susceptibility SNPs or the *polygenicity*. The Gaussian slab is parameterized with mean 0 and variance 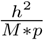 if the genotypes are standardized and phenotypes have a variance of 1, where *h*^2^ is the heritability of the trait. To facilitate the explanation, we denote the effect sizes for susceptibility SNPs as the (*M* × 1) vector ***γ*** = (*γ*_1_, …, *γ*_*M*_)^*T*^. If SNP *m* is a susceptibility SNP, the true effect size *β*_*m*_ will be *γ*_*m*_ and 0 otherwise according to its susceptibility status *c*_*m*_:

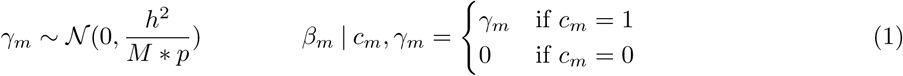

We can model the conditional distribution of the GWAS effect sizes given the true effect sizes, where 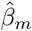 is the estimated marginal effect size of the *m*^*th*^ SNP from GWAS and the (*M* × 1) vector 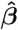 represents the estimated effect sizes for all SNPs. The covariance matrix is parameterized by the environmental error and the (*M* × *M*) LD matrix, which can be computed using genotypes from the study, 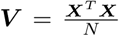. Here we parameterize the environmental error as 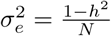 :

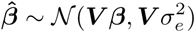

We next impose a symmetric beta prior on the polygenicity parameter, *p*. In practice, we set *θ*_*p*_ = 0.2, but find that our model is robust to the choice of other hyper-parameter choices (see Section 3.2).

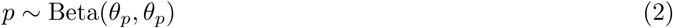

Finally, the joint distribution is given by:

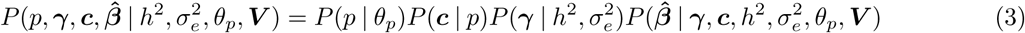

### 2.2 Parameter inference in our model

The true posterior distribution of each parameter is intractable; thus, we use Markov Chain Monte Carlo (MCMC) to approximate the posterior distribution of each parameter. Specifically, we derive a Gibbs sampler [10] to sample from the posterior distribution for the polygenicity parameter *p* and latent variables ***c, γ***. In this work, we only focus on accurately estimating the polygenicity *p* since other quantities such as the heritability can be estimated using previous methods [5] and then used as an input to our model. Although our framework estimates the SNP effect sizes and susceptibility statuses (***c, γ***), we only report the posterior distribution of *p* since we are interested in estimating only the polygenicity.

The generative model and inference for estimating genome-wide polygenicity and regional polygenicity are identical under the assumption that there is no correlation between SNPs across different regions. Note that the regional setting still assumes correlations between SNPs within regions. Because LD tends to diminish with genomic distance, we expect this to be the case for most regions in the genome. Therefore, we can use our defined model of genome-wide polygenicity and directly apply it in the regional setting under this set of assumptions.

To summarize, as input, our method takes in marginal effect sizes from GWAS for a single trait for a given region 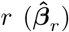, LD (***V*** _*r*_) for region *r*, an estimate of the SNP heritability per region (*h*_*r*_^2^), and the sample size of the GWAS study (*N*). As output, BEAVR estimates the posterior distribution of the regional polygenicity (*p*_*r*_). Note that for clarity, we drop the per region notation and refer to the regional heritability and regional polygenicity as *h*^2^ and *p*, and LD matrix and GWAS effects for the region as ***V*** and 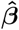 since the model and inference procedure do not change between the regional and genome-wide setting. Furthermore, any parameters that refer to genome-wide quantities will be explicitly noted.

### 2.3 Gibbs sampler

Given the formation of the joint posterior probability, we can approximate this distribution through Gibbs sampling, a type of MCMC algorithm that generates samples from the conditional distribution of each parameter.

#### Transforming GWAS effect sizes

First, we transform the marginal effects from GWAS by multiplying by 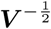 to de-correlate the effects in the covariance term, which provides a diagonal covariance matrix that lends itself to more straightforward derivations of conjugate distributions. However, we note that the effect sizes are not fully de-correlated, as LD still affects the mean of the effect size distribution. We refer to these transformed GWAS effect sizes as 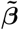 and use these throughout the rest of the derivations. The distribution for the transformed effect sizes from GWAS becomes:

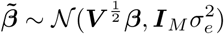

Here, ***I***_*M*_ is the identity matrix of size *M* × *M*. We note that this is a one-time transformation that is performed before running the sampler. These transformed effects can be efficiently stored and computed for each genomic region beforehand.

#### Sampling *γ* and *c*

First, we recall that the true effect sizes at each SNP (*β*_*m*_) assume a spike-and-slab prior parameterized by the susceptibility SNP effect size (*γ*_*m*_) and the susceptibility status (*c*_*m*_) (see Equation 1). Because the susceptibility effect size and status are tightly correlated (i.e. *γ*_*m*_ = 0 if *c*_*m*_ = 0 and *γ*_*m*_ ≠ 0 if *c*_*m*_ = 1), we choose to sample both *γ*_*m*_ and *c*_*m*_ together in a block step.

Let 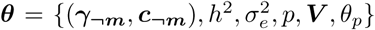, where ***γ***_***¬m***_ denotes all susceptibility effect sizes except for the effect of the *m*^*th*^ SNP; this similarly follows for ***c***_***¬m***_.

By the chain rule, we have:

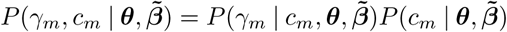

We are interested in the posterior distribution of the susceptibility effect size *γ*_*m*_ when *c*_*m*_ = 1 since *P* (*γ*_*m*_ | *c*_*m*_ = 0) = 0 due to the spike-and-slab prior. This can be expressed as:

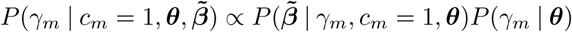

Working with the transformed GWAS effect sizes and the properties of the conjugate priors, the posterior distribution of *γ*_*m*_ becomes univariate Gaussian with the following mean and variance. Here we denote 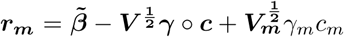, which can be thought of as the residual from subtracting the effects of all SNPs except for SNP *m*. We define 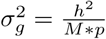 and 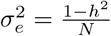 which stems from the generative model defined in Section 2.1. The full derivation can be found in the Supplementary Materials.

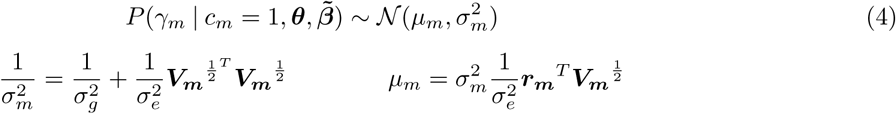

We must also determine the susceptibility status of each SNP at each iteration. This is computed by sampling 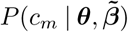:

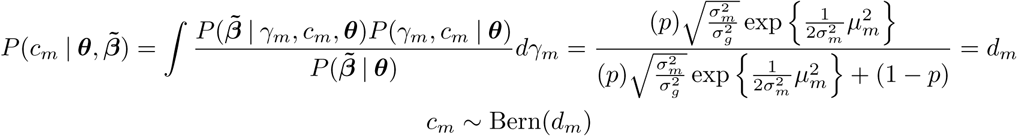

#### Sampling *p*

The conditional posterior distribution of *p* depends not only on the susceptibility status of each SNP (*c*_*m*_), but also on the susceptibility SNP effect sizes (*γ*_*m*_) since *p* parameterizes the variance term of *γ*_*m*_. Thus, we are not able to leverage conjugacy to write a closed form for the conditional posterior distribution. Instead, we can sample from the distribution using a random-walk Metropolis-Hastings step [11].

This step generates a sample *p*^(*t*)^ at iteration *t* given the sample *p*^(*t*-1)^ from the previous iteration using the chosen proposal distribution, *Q*(*p*^*^ | *p*^(*t*-1)^), which generates a proposed sample *p*^*^. This sample is then either accepted or rejected depending on the Metropolis-Hastings ratio which depends on the ratio of the posterior probability density at the proposed parameter compared to the density at the parameter from the previous iteration. We define our proposal distribution as follows:

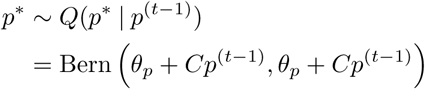

Here, *C* is a constant that controls the variance of the proposal distribution. In practice we found that *C* = 10 yields effective mixing.

The final step in specifying the Metropolis-Hastings algorithm lies in computing the ratio of the posterior probability density at the proposed parameter (*p*^*^) to the previous parameter (*p*^(*t*-1)^). Computing this ratio only requires writing the posterior distribution up to a normalization constant:

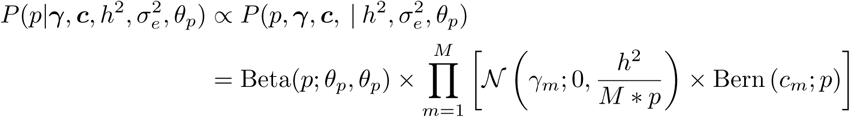

### 2.4 Reducing complexity from 𝒪(*M* ^2^) to 𝒪(*MK*)

The key computational bottleneck in estimating the regional polygenicity from our Gibbs sampling scheme is calculating the mean of the posterior for the susceptibility SNPs’ effect sizes (*µ*_*m*_) in Equation 4. Calculating the residual term, 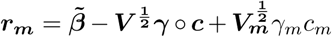, is 𝒪(*M*) because we must subtract the effects of all SNPs in the region due to LD. However, this computation must be performed for every SNP at each iteration, making the final complexity of one iteration of the sampler to be 𝒪(*M* ^2^) and thus too computationally costly for regions with more than a few SNPs. However, we expect most complex traits to have a small proportion of susceptibility SNPs. Formally, we denote the total number of susceptibility SNPs as *K* and *M* as the total number of SNPs in the region. As the sampler converges to the stationary distribution, we would expect the vector of susceptibility statuses ***c*** to be very sparse, where *K* ≪ *M*.

By expanding Equation 4 and re-writing ***r***_***m***_ in summation notation, we can reconfigure this update to only depend on *K* when sampling each SNP. This value can be efficiently computed in each MCMC iteration and calculated using the updating scheme below:

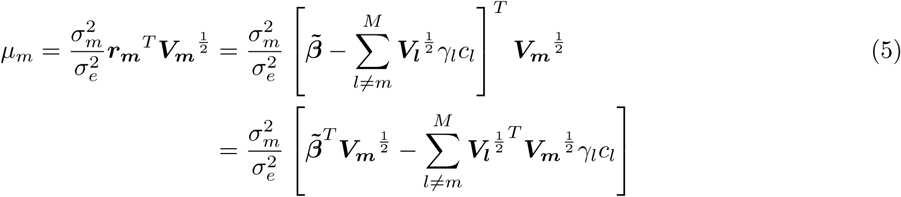

Importantly, the term 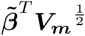 is only constructed from GWAS marginal effect sizes and the LD matrix, which does not change throughout the sampler. Thus, we can construct 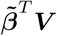 only one time and then simply access the *m*^*th*^ element of this vector when sampling each SNP. A similar step can be performed for the 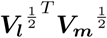 term where we only initially compute 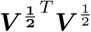 and then access the (*l, m*) element at each step. The total operation in Equation 4 reduces to a very efficient vector subtraction where we only consider SNPs where *c*_*l*_ = 1 since we do not need to consider elements where *c*_*l*_ = 0 and thus *γ*_*l*_ = 0. After performing this calculation for each SNP, the overall run time for each iteration of the sampler becomes 𝒪(*MK*). The full algorithm is further detailed in Algorithm 1.

#### Algorithm 1

Gibbs sampler

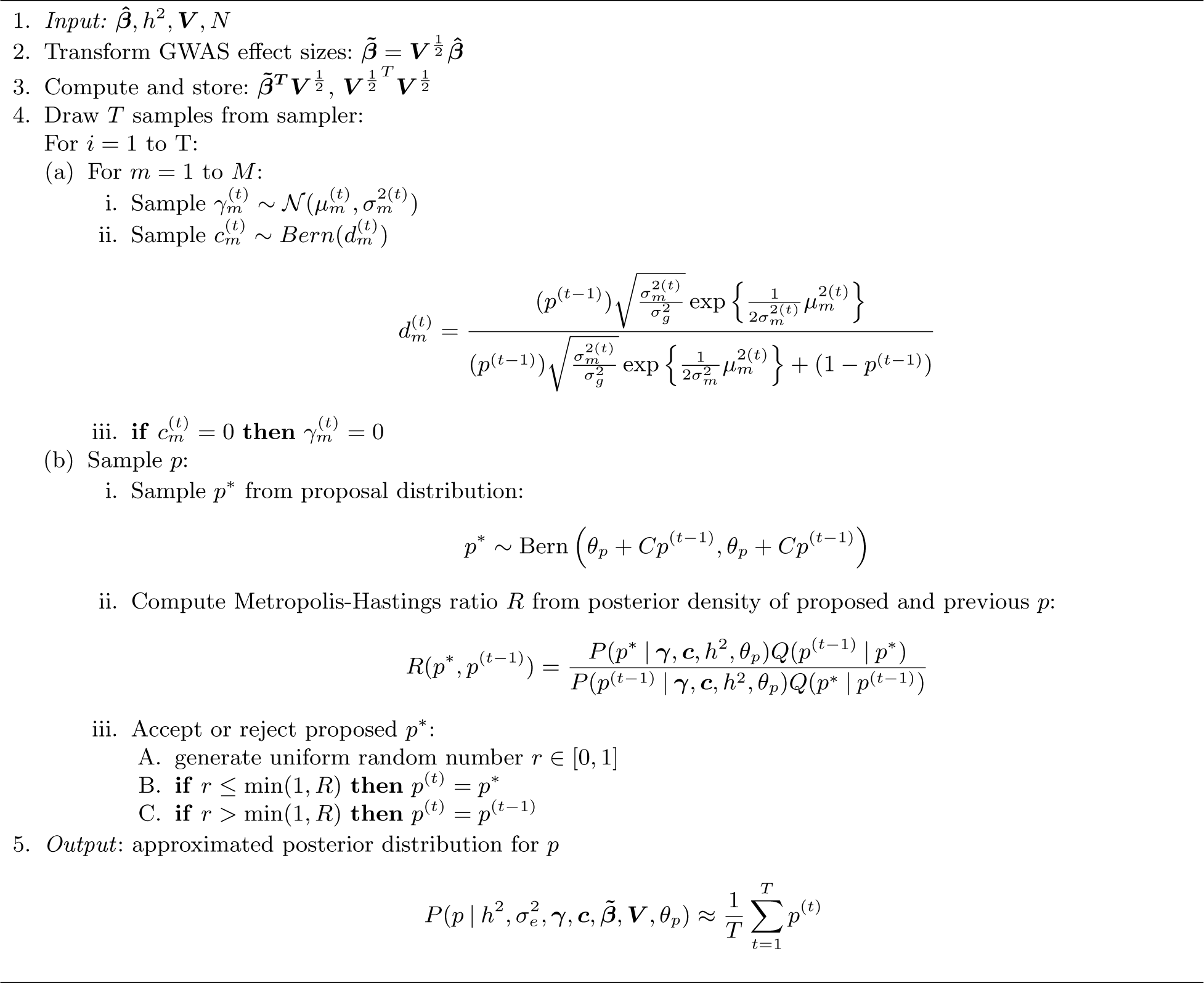

## 3 Results

### 3.1 BEAVR fully models correlation between SNPs

To our knowledge, the only published method that can estimate polygenicity from GWAS effects is GENESIS [4], which uses a mixture model to estimate the number of susceptibility SNPs for a trait. GENESIS employs a mixture of either 2 or 3 mixture components (either 1 or 2 normal distributions and a point mass) to capture both large and small effect sizes from susceptibility SNPs. However, GENESIS was developed to do genome-wide scale analyses and accommodates this by making the assumption that LD patterns are independent of the probability of a SNP belonging to different mixture components. By way of this assumption, the estimation of polygenicity does not require the full conditioning of every SNP in the genome and thus does not require fully modeling all correlations between SNPs. However, we hypothesize that this approximation will lead to a downward bias. Because LD spreads the effects of SNPs throughout a region, not considering the full LD structure will limit the method’s ability to fully recapture all susceptibility SNPs. Thus, methods such as GENESIS that make this assumption are likely not able to provide the fine-grain resolution necessary for analyses at a regional level. To accommodate the large computational bottleneck induced by LD, we derive a computationally efficient inference algorithm that makes no simplifying assumptions when modeling LD. Instead, we leverage the expected sparsity of complex traits and at each iteration of the algorithm, we only consider the sampled susceptibility SNPs when estimating polygenicity.

### 3.2 Simulations

We generated marginal effect sizes for a region of *M* = 1000 SNPs from a synthetic GWAS and assume that the region is selected from a GWAS with 500K SNPs genome-wide, which corresponds to the typical number of SNPs on a genotyping array. To simulate susceptibility SNPs, we drew the susceptibility status of each SNP based on the polygenicity parameter where *c*_*m*_ = 1 denotes a susceptibility SNP and *c*_*m*_ = 0 otherwise. Then, if *c*_*m*_ = 1, the effect size of SNP *m* was drawn from a univariate Gaussian distribution with mean 0 and variance equal to the genome-wide heritability 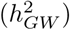 divided by the number of array SNPs multiplied by the polygenicity. By using the total number of array SNPs in the denominator, we can use genome-wide heritability values when simulating, which is typically more interpretable.

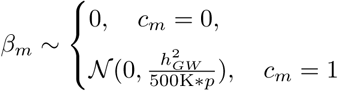

To compute the marginal effect size of each SNP estimated from GWAS, 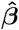, we draw the estimated effect sizes from a conditional distribution of the GWAS effect sizes, as described in Section 2.1. We use LD computed from a region with 1000 SNPs from chromosome 22 from 337K individuals from the UK Biobank as our correlation matrix ***V***.

First, we confirm that our method accurately predicts regional polygenicity under a variety of genetic architectures and sample sizes. We consider different values for the regional polygenicity to be 0.005, 0.01, 0.05, and 0.10. In the first scenario, we fix the genome-wide heritability to be 0.50 and the sample size to be 500K individuals, which is comparable to the sample sizes of many current GWAS studies [12, 13].

Each experiment is repeated 100 times in order to assess the variance of each method. We ran BEAVR for 1000 iterations and used the first 250 iterations as burn-in. We used the same LD information that was used for simulation (*i.e.* “in-sample” LD). For the heritability parameter, we use the simulated genome-wide heritability scaled by the number of SNPs in the region and the number of SNPs on the array 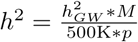. We ran GENESIS using the default parameter settings and LD information from 1000 Genomes [14]. We use both the 2-component and 3-component settings when running GENESIS. We note that GENESIS was designed to be run with external LD information and that providing in-sample LD information was not a utility of the current implementation of the method.

As shown in Figure 1, we can see that BEAVR performs robustly across each scenario. However, in settings with higher polygenicity, GENESIS demonstrates a severe downward bias. This observation supports our initial hypothesis that not modeling the full LD structure limits the ability of GENESIS to fully recover all susceptibility SNPs. GENESIS cannot recapture the effects of SNPs with smaller effect sizes that have been spread throughout the region due to LD. Although GENESIS was run with external LD information, the authors of the method also observe a downward bias in estimates of *p* in their analyses, leading us to believe that this observed bias is not an artifact of external LD.

**Fig. 1.**
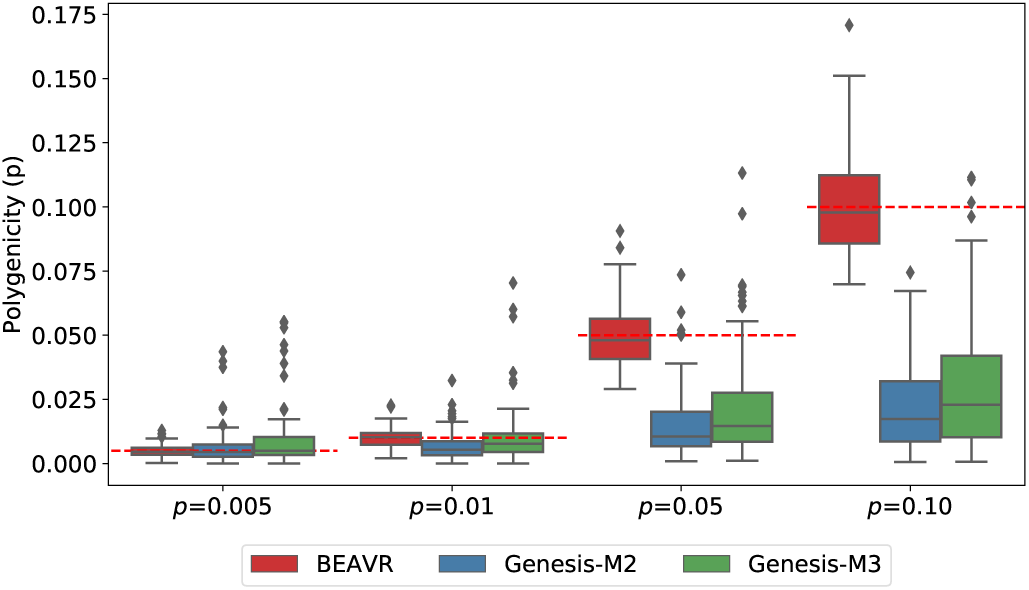
**BEAVR is relatively unbiased in simulated data:** We simulated 100 regions where the genome-wide heritability was set to 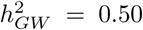 and the true polygenicity of the region was *p* = 0.005, 0.01, 0.05, 0.10. We compared BEAVR to GENESIS-M2 (2-component) and GENESIS-M3 (3-component). The x-axis denotes the simulated values for polygenicity and the y-axis denotes the estimated values across 100 replicates. Dashed red lines denote the true polygenicity value in each setting.

We perform additional experiments where we vary the genome-wide heritability to be 0.10 and 0.25 and the sample size to be 50K and 1 million individuals (Supplementary Figure 1) to fully explore the limitations of our method. We note that when sample sizes are small (*N* = 50*K* individuals), BEAVR starts to demonstrate a downward bias when *p* ≥ 0.05. This is because the environmental noise caused by the low sample size is a similar magnitude as the effect sizes from susceptibility SNPs, making it difficult to correctly identify the susceptibility status of a SNP. However, most GWAS studies have sample sizes in the hundreds of thousands and more recent studies even having a million samples; therefore, we do not expect this to be a limitation in the utility of our method and recommend only applying BEAVR on well-powered studies. Second, we observe a slight downward bias when the genome-wide heritability is very low 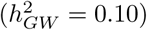 and the polygenicity is high (*p* ≥ 0.05). When the heritability is spread across many SNPs, the effect sizes become very close to zero, leading to the observed downward bias. However, this bias is only moderate when the sample size is high (N=1 million individuals). Therefore, we advise using BEAVR on traits with 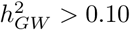 or when sample sizes are large enough (*N* ≥ 1 million individuals).

We perform an additional experiment that mirrors exactly how BEAVR would be applied in practice. First, we simulate phenotypes using genotype information from chromosome 22 (M=9564 SNPs) for N=337K individuals from the UK Biobank. Each phenotype is simulated to have 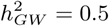 and polygenicity *p* = 0.01. When simulating the effect sizes of susceptibility SNPs, we couple the magnitude of a SNP’s effect size with the SNP’s minor allele frequency, which is believed to be a more biologically accurate genetic architecture [15]. The simulated effect sizes follow the framework outlined in previous work[16]. We then perform a marginal analysis to compute the estimated effect sizes. Second, we divide the simulated data into consecutive regions of 6Mb for a total of 6 regions, where on average there are 1000 SNPs in each block. Third, instead of using known values for the regional heritability, we use the HESS software [5]– a software that estimates the heritability within a region from GWAS marginal effects– to estimate this quantity. We find that our estimates of polygenicity are unbiased across all regions as shown in Figure 2(a). Although there exists LD that spans between regions, leading to correlation between SNPs across regions, we did not find this to affect the accuracy of our results. This is likely because the magnitude of correlation due to LD diminishes with genomic distance.

**Fig. 2.**
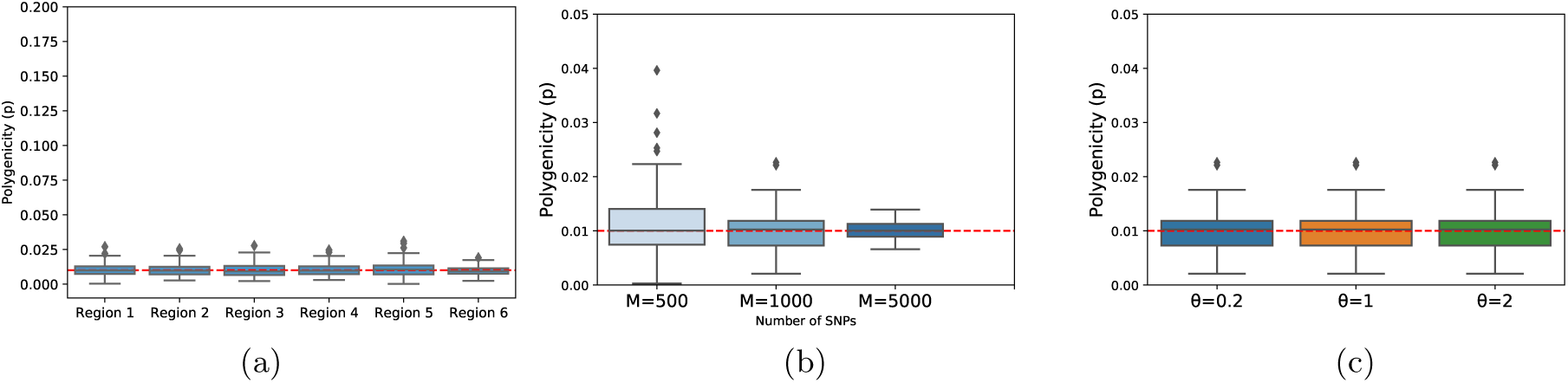
**(a) BEAVR is unbiased in realistic study settings:** Using SNP data from chromosome 22 (M=9564 array SNPs, N=337K individuals), we simulated 100 replicates where the genome-wide heritability was 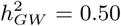 and *p* = 0.01. We divided the data into 6Mb consecutive regions for a total of 6 regions and estimated the regional heritabilities using HESS. Using BEAVR and the estimated heritabilities, we estimated the regional polygenicities to be unbiased across all regions. **(b) BEAVR is robust across various region sizes:** We ran 100 replicates where the genome-wide heritability is fixed 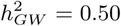, polygenicity *p* = 0.01, sample size *N* = 500*K*, and varied the number of SNPs in the region from *M* = 500, 1*K*, 5*K* SNPs. We used BEAVR to estimate the polygenicity in each region and found our results to be unbiased across all regions. **(c) BEAVR is robust to different priors:** We ran 100 replicates where the genome-wide heritability was set to 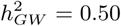, local polygenicity *p* = 0.01, and a sample size of *N* = 500*K*. We find that the accuracy of our results is invariant to our choice of prior hyper-parameters.

We also vary the robustness of our model across different sized regions. Using a simulated GWAS with genome-wide heritability 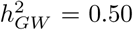, sample size *N* = 500*K*, and polygenicity *p* = 0.01, we vary the size of the region from *M* = 500, 1*K*, 5*K* SNPs. From Figure 2(b), we can see that our estimates are robust across regions of various sizes and demonstrates the potential utility of applying BEAVR across different sized regions for a variety of biological questions, such as estimating the regional polygenicity around genes or within larger LD blocks.

Finally, we assess the robustness of our results when using different hyper-parameters for our prior. Using a simulated GWAS with genome-wide heritability 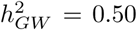, sample size *N* = 500*K*, and polygenicity *p* = 0.01 we vary our choice of hyper-parameters for our prior: *θ*_*p*_ = 0.2, 1, 2. We find that the accuracy of our results does not heavily depend on our choice of prior, as shown in Figure 2(c), and we use continue to use *θ*_*p*_ = 0.2 for all subsequent analyses.

### 3.3 Computational efficiency

We next assess the computational efficiency of our method. We compared two implementations of BEAVR, one with the algorithmic speedup and one without. The algorithmic speedup as described in Section 2.4 only considers SNPs that are sampled as susceptibility SNPs at that iteration. This reduces the original implementation of 𝒪(*M* ^2^), where M is the number of SNPs, to 𝒪(*MK*), where K is the number of susceptibility SNPs. Since regional analyses can include regions of the genome that vary in size, we wanted to assess the efficiency of our method across larger regions of the genome. In the first experiment, we simulated GWAS data for M=500, 1K, 2K, 3K, 4K, 5K SNPs with a regional polygenicity of 0.01. We generated 100 replicates and computed the average time per iteration of the MCMC sampler. All comparisons were done on an Intel(R) Xeon(R) CPU 2.10 GHz server with 128GB RAM and run on a single core. From Figure 3(a), we see that the runtime of BEAVR (Speedup) is approximately linear in the number of SNPs, compared with the baseline implementation which is quadratic in the number of SNPs.

**Fig. 3.**
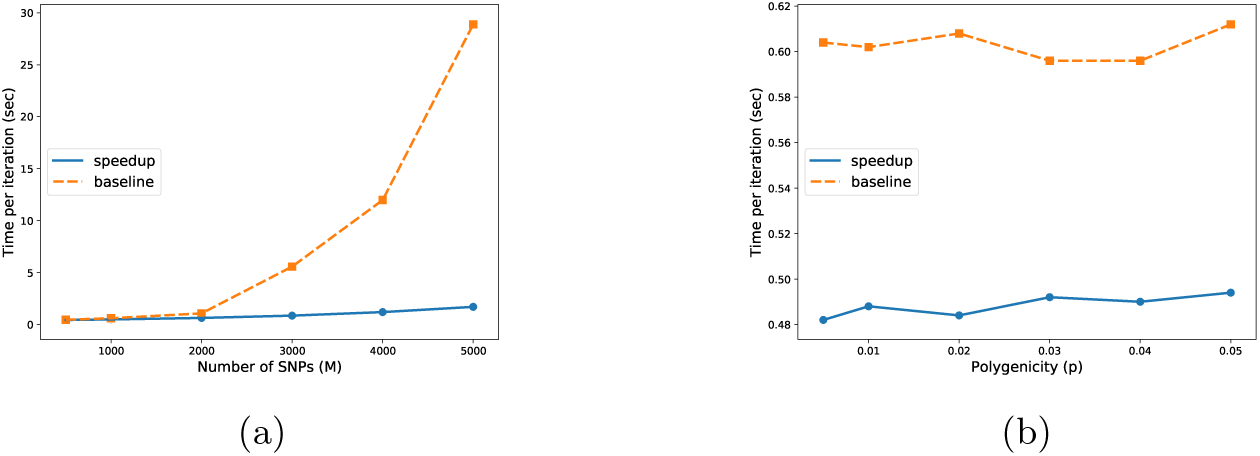
**(a) BEAVR is efficient across different sized regions:** We compare the version of BEAVR with the algorithmic speedup (‘speedup’) versus the implementation without any speedup (‘baseline’). For 100 replicates, we vary the number of SNPs in the region while fixing the polygenicity of each region to *p* = 0.01. We report the runtime of each method in terms of average number of seconds per iteration of the MCMC sampler. **(b) BEAVR is efficient across varying levels of polygenicity:** We fix the number of SNPs in the region to be *M* = 1000 while varying the polygenicity. Although the theoretical complexity of our algorithm is 𝒪(*MK*), in practice, the run-time of BEAVR is closer to 𝒪(*M*) since complex traits tend to have *K* ≪ *M*.

Since the runtime of the algorithm depends on *K*, the number of susceptibility SNPs, we perform additional experiments to assess the role of this value in the efficiency of our algorithm. We simulated GWAS data for 1000 SNPs and vary the regional polygenicity from p=0.005, 0.01, 0.02, 0.03, 0.04, 0.05. Figure 3(b) shows that the efficiency of our method is robust to the regional polygenicity, lending itself to traits of varying genetic architectures.

### 3.4 Empirical Analysis of BMI, eczema, and high cholesterol

We computed marginal effect sizes for BMI, eczema, and high cholesterol from the UK Biobank [8]. We restrict our analyses to a subset of unrelated individuals identified as ‘White British’ (N = 290K individuals, M=460K array SNPs). We divide the genome into a total of 470 6Mb regions where each region has on average 1000 SNPs. For each region, we use HESS to compute an estimate of the regional heritability. We then use BEAVR to estimate the regional polygenicities of all regions across the genome for each of the three traits. We report a nonzero regional polygenicity if the posterior mean is > 2 standard deviations away from 0.

From Figure 4(a), we can see that all traits have many susceptibility SNPs spread throughout the genome, which is consistent with previous findings [6]. However, we note that BMI has susceptibility SNPs in almost every region of the genome, whereas eczema and high cholesterol show more sparse architectures. From Figure 4(b), the histogram of regional polygenicities for BMI shows that the trait is mostly influenced by regions with many susceptibility SNPs. This contrasts with eczema which has mostly regions with only a small percentage of susceptibility SNPs. Overall, this result is consistent with previous findings that show that BMI has overall more susceptibility SNPs than autoimmune traits such as eczema [6], but provides evidence that even though eczema has fewer susceptibility SNPs, these SNPs are distributed approximately uniformly throughout the genome.

**Fig. 4.**
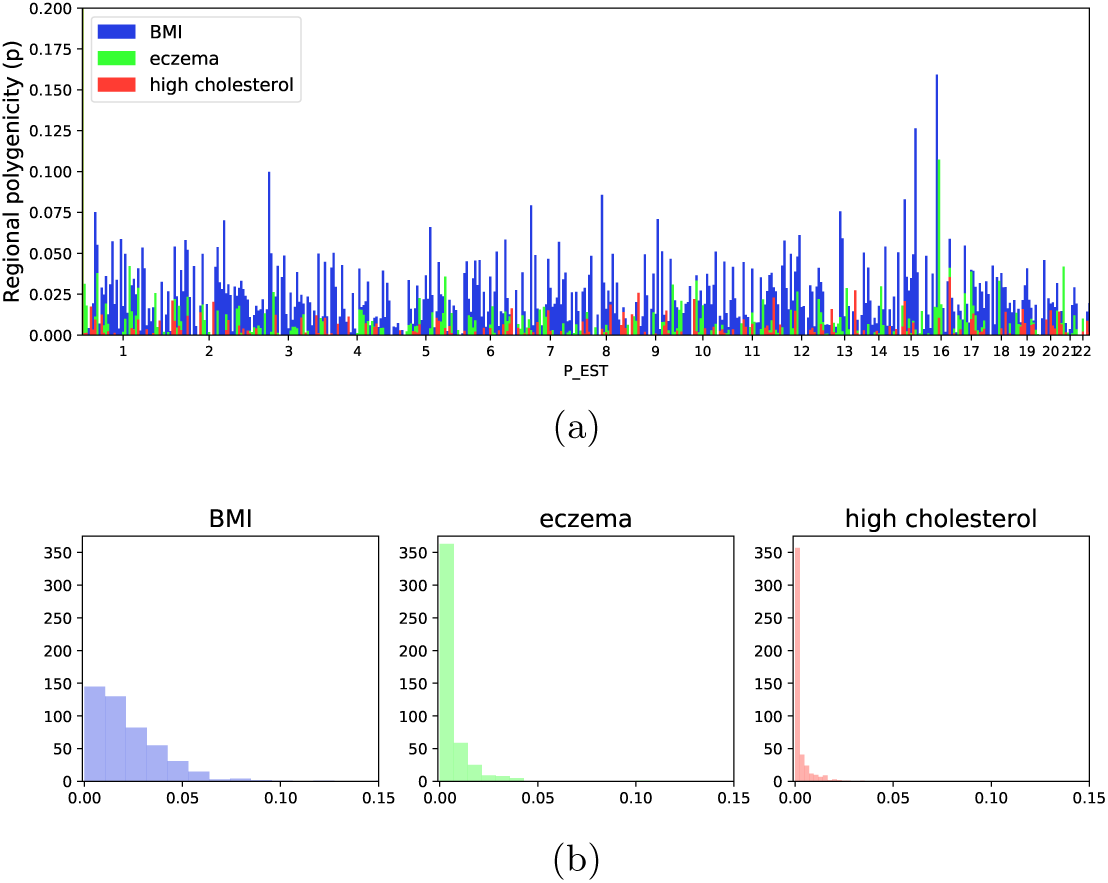
**Manhattan-style plot of regional polygenicity across the genome:** We divide the genome into 6Mb regions and estimate the regional polygenicity for each region for BMI, eczema, and high cholesterol. We plot the regional polygenicity for each region across the genome versus the genomic position of the region. The x-axis denotes the corresponding chromosome. **Histogram of regional polygenicity for BMI, eczema, and high cholesterol:** We visualize the distribution of regional polygenicities for each trait with a histogram of our estimates. BMI has many regions with a high fraction of susceptibility SNPs whereas eczema and high cholesterol are mostly described by many regions with only a few susceptibility SNPs in each region.

## 4 Discussion

We propose BEAVR, a novel, scalable method to estimate regional polygenicity that models the full correlation structure among SNPs in a Bayesian framework. We employ a fast inference procedure that leverages the assumption that susceptibility SNPs for complex traits compose a small fraction of the genome, achieving an almost linear runtime, 𝒪(*MK*), for *M* SNPs and *K* susceptibility SNPs. We show that our method achieves higher accuracy than methods that only model first-order correlations among SNPs and is robust across a variety of genetic architectures. Using BEAVR, we partition polygenicity by genomic regions and show how the distribution of regional polygenicity varies across complex traits.

We conclude by discussing limitations of our study and future directions. First, while our approach can be applied to GWAS summary statistics (marginal regression coefficients) given an estimate of LD, which would typically be obtained from a reference panel, in this work, we apply BEAVR using only in-sample LD. The robustness of BEAVR to reference panel LD will be investigated in future work. Second, we want to clearly distinguish BEAVR from methods that perform statistical fine-mapping. In contrast to BEAVR, which reports the proportion of susceptibility SNPs in a region, fine-mapping aims to prioritize susceptibility SNPs within a region for follow-up studies [17]. Thus, while related to regional polygenicity, fine-mapping approaches output the posterior probability of each SNP to be a susceptibility SNP and are typically applied only to GWAS significant regions where we are confident that there is at least one susceptibility SNP driving the association signal. In addition, many publicly available GWAS summary statistics were computed using linear mixed models (LMMs) rather than ordinary least squares to control for population structure. Association statistics computed from LMMs have different statistical properties than those from ordinary least squares; we leave an investigation of BEAVR with mixed model association statistics to future work.

## 5 Code availability

The code for our method is freely available: https://github.com/bogdanlab/BEAVR We additionally provide a full tutorial with test data.

## 6 Acknowledgements

We are grateful to Yi Ding, Tommer Schwarz, and Alec Chiu for their helpful and insightful discussions. This research was conducted using the UK Biobank Resource under applications 33297 and 33127. This work was funded in part by National Institutes of Health (NIH), under awards R01HG009120, R01HG006399, U01CA194393, R00GM111744, R35GM125055, National Science Foundation Grant III-1705121, an Alfred P. Sloan Research Fellowship, and a gift from the Okawa Foundation.

