## Supplemental Materials for "A scalable method for estimating the regional polygenicity of complex traits"

### Additional simulations

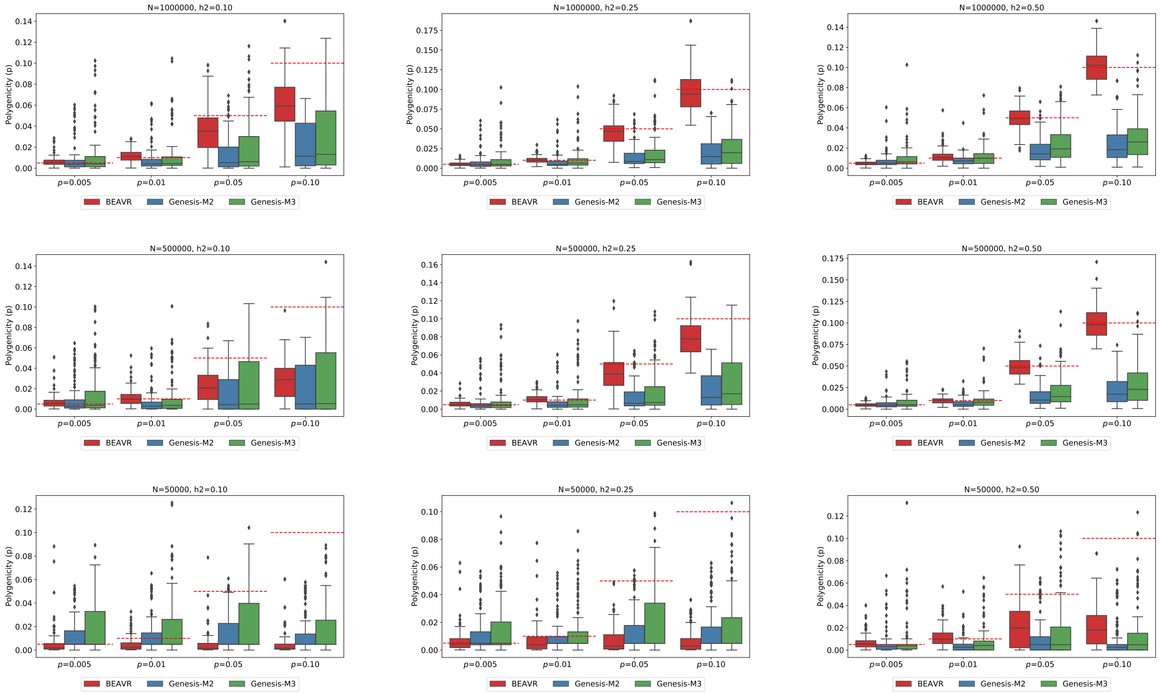


**Fig. 1. BEAVR is relatively unbiased in simulated data**: We ran 100 replicates where we vary the genome-wide heritability to be $h^2_{GW} = 0.10, 0.25, 0.5$, the polygenicity of the region to be $p = 0.005, 0.01, 0.05, 0.10$, and the sample size N=50K, 500K, 1 million individuals. We compared BEAVR to GENESIS-M2 (2-component) and GENESIS-M3 (3-component). The x-axis denotes the simulated values for polygenicity and the y-axis denotes the estimated values across 100 replicates. Dashed red lines denote the true polygenicity value in each setting.

### Additional derivations

#### Sampling $\boldsymbol{\gamma_m}, \mathbf{c_m}$

We derive a Gibbs sampler to sample from the posterior distribution of each parameter $\{c_m, \gamma_m, p\}$. Because a SNP's causal status and effect size are highly correlated, we sample $(\gamma_m, c_m)$ together in a block.

Let $\theta = \{(\boldsymbol{\gamma_{\neg m}}, \boldsymbol{c_{\neg m}}), \sigma_g^2, p\}$, where $\boldsymbol{\gamma_{\neg m}}$ denotes all effect sizes except for the effect of the $m^{th}$ SNP; this similarly follows for $\boldsymbol{c_{\neg m}}$. We denote $\sigma_g^2 = \frac{h^2}{M*p}$ and $\sigma_e^2 = \frac{1-h^2}{N}$. The derivation for each marginal posterior distribution, $P(\gamma_m \mid \cdot)$ and $P(c_m \mid \cdot)$ is given separately below. By the chain rule note that:

$$P(\gamma_m, c_m \mid \theta, \tilde{\boldsymbol{\beta}}) = P(\gamma_m \mid c_m, \theta, \tilde{\boldsymbol{\beta}})P(c_m \mid \theta, \tilde{\boldsymbol{\beta}})$$

Additionally, for convenience we'll denote $\boldsymbol{r_m} = \tilde{\boldsymbol{\beta}} - \boldsymbol{V}^{\frac{1}{2}}\boldsymbol{\gamma c} + \boldsymbol{V_m^{\frac{1}{2}}}\gamma_m c_m$, which can be thought of the residual from subtracting the effects of all SNPs except for SNP $m$.

Deriving the first term, $P(\gamma_m \mid c_m, \theta, \tilde{\boldsymbol{\beta}})$, we'll break up the expression into the cases when $c_m = 1$ and $c_m = 0$:

$$\begin{aligned}
P(\gamma_m \mid c_m = 1, \theta, \tilde{\boldsymbol{\beta}}) &\propto P(\tilde{\boldsymbol{\beta}} \mid \gamma_m, c_m = 1, \theta)P(\gamma_m \mid \theta) \\
&= \exp\Big\{ -\frac{1}{2\sigma_e^2}(\tilde{\boldsymbol{\beta}} - \boldsymbol{r_m})^T(\tilde{\boldsymbol{\beta}} - \boldsymbol{r_m})\Big\}\exp\Big\{ -\frac{1}{2\sigma_g^2}\gamma_m^2\Big\} \\
&= \exp\Big\{ -\frac{1}{2\sigma_e^2}(\boldsymbol{r}^T\boldsymbol{r} - 2\boldsymbol{r_m}^T\boldsymbol{V_m}^{\frac{1}{2}}\gamma_m + \boldsymbol{V}_m^{\frac{1}{2}^T}\boldsymbol{V}_m^{\frac{1}{2}}\gamma_m^2) + -\frac{1}{2\sigma_g^2}\gamma_m^2\Big\} \\
&\text{[drop constants that don't depend on } \gamma_m] \\
&= \exp\Big\{ -\frac{1}{2\sigma_e^2}(-2\boldsymbol{r_m}^T\boldsymbol{V_m}^{\frac{1}{2}}\gamma_m + \boldsymbol{V}_m^{\frac{1}{2}^T}\boldsymbol{V}_m^{\frac{1}{2}}\gamma_m^2) + -\frac{1}{2\sigma_g^2}\gamma_m^2\Big\} \\
&\text{[common denominators]} \\
&= \exp\Big\{ -\frac{1}{2\sigma_e^2 2\sigma_g^2}(-2\boldsymbol{r}_m^T\boldsymbol{V_m}^{\frac{1}{2}}2\sigma_g^2\gamma_m + \boldsymbol{V_m}^{\frac{1}{2}^T}\boldsymbol{V}_m^{\frac{1}{2}}2\sigma_g^2\gamma_m^2 + 2\sigma_e^2\gamma_m^2)\Big\} \\
&= \exp\Big\{ -\frac{\gamma_m^2}{2}\left(\frac{1}{\sigma_g^2} + \frac{1}{\sigma_e^2}\boldsymbol{V}_m^{\frac{1}{2}^T}\boldsymbol{V}_m^{\frac{1}{2}}\right) + \gamma_m\left(\frac{1}{\sigma_e^2}\boldsymbol{r}_m^T\boldsymbol{V}_m^{\frac{1}{2}}\right)\Big\} \\
&= \exp\Big\{ -\frac{\gamma_m^2}{2}(a) + \gamma_m(b)\Big\} \\
&\Big(a = -\frac{1}{2\sigma_m^2}, b = \frac{\mu_m}{\sigma_m^2}\Big) \\
&= \mathcal{N}(\mu_m, \sigma_m^2) \\
&\frac{1}{\sigma_m^2} = \frac{1}{\sigma_g^2} + \frac{1}{\sigma_e^2}\boldsymbol{V_m}^{\frac{1}{2}^T}\boldsymbol{V_m}^{\frac{1}{2}} \\
&\mu_m = \sigma_m^2\frac{1}{\sigma_e^2}\boldsymbol{r_m}^T\boldsymbol{V_m}^{\frac{1}{2}}
\end{aligned}$$

$$P(\gamma_m \mid c_m = 0, \theta, \tilde{\boldsymbol{\beta}}) = 0$$

The bottom line follows because if a SNP is non-causal ($c_m = 0$), the effect size is 0.

Deriving the second term, $P(c_m \mid \theta, \tilde{\boldsymbol{\beta}})$:

$$
\begin{aligned}
P(c_m \mid \theta, \tilde{\boldsymbol{\beta}}) &= \int P(c_m, \gamma_m \mid \theta, \tilde{\boldsymbol{\beta}}) d\gamma_m \\
&= \int \frac{P(\tilde{\boldsymbol{\beta}} \mid \gamma_m, c_m, \theta) P(\gamma_m, c_m \mid \theta)}{P(\tilde{\boldsymbol{\beta}} \mid \theta)} d\gamma_m \\
&= \int \frac{P(\tilde{\boldsymbol{\beta}} \mid \gamma_m, c_m, \theta) P(\gamma_m \mid c_m \theta) P(c_m \mid \theta)}{P(\tilde{\boldsymbol{\beta}} \mid \theta)} d\gamma_m \\
&= \frac{P(c_m \mid \theta)}{P(\tilde{\boldsymbol{\beta}} \mid \theta)} \int P(\tilde{\boldsymbol{\beta}} \mid \gamma_m, c_m, \theta) P(\gamma_m \mid c_m, \theta) d\gamma_m \\
&\text{[denominator does not depend on } c_m\text{]} \\
&= P(c_m \mid \theta) \int \Big[ \frac{1}{\sqrt{2\pi\sigma_m^2}} \exp\Big\{ -\frac{1}{2\sigma_m^2} (\gamma_m - \mu_m)^2 \Big\} \Big] d\gamma_m \sqrt{2\pi\sigma_m^2} \exp\Big\{ -\frac{1}{2\sigma_e^2} \boldsymbol{r_m}^T \boldsymbol{r_m} + \frac{1}{2\sigma_m^2} \mu_m^2 \Big\} \\
&= P(c_m \mid \theta) \frac{\sqrt{2\pi\sigma_m^2}}{\sqrt{2\pi\sigma_e^2}\sqrt{2\pi\sigma_g^2}} \exp\Big\{ -\frac{1}{2} \frac{\boldsymbol{r_m}^T \boldsymbol{r_m}}{\sigma_e^2} + \frac{1}{2\sigma_m^2} \mu_m^2 \Big\}
\end{aligned}
$$

To sample $c_m$, we'll draw $c_m \sim \text{Bern}(d_m)$, where $d_m$ is defined as follows:

$$
\begin{aligned}
d_m &= \frac{\frac{\sqrt{2\pi\sigma_m^2}}{\sqrt{2\pi\sigma_e^2}\sqrt{2\pi\sigma_g^2}} P(c_m = 1) \exp\Big\{ -\frac{1}{2} \frac{\boldsymbol{r}_m^T \boldsymbol{r}_m}{\sigma_e^2} + \frac{1}{2\sigma_m^2} \mu_m^2 \Big\}}{\frac{\sqrt{2\pi\sigma_m^2}}{\sqrt{2\pi\sigma_e^2}\sqrt{2\pi\sigma_g^2}} P(c_m = 1 \mid \theta) \exp\Big\{ -\frac{1}{2} \frac{\boldsymbol{r}_m^T \boldsymbol{r}_m}{\sigma_e^2} + \frac{1}{2\sigma_m^2} \mu_m^2 \Big\} + \frac{1}{\sqrt{2\pi\sigma_e^2}} P(c_m = 0) \exp\Big\{ -\frac{1}{2\sigma_e^2} \boldsymbol{r}_m^T \boldsymbol{r}_m^T \Big\}} \\
&= \frac{\frac{\sqrt{2\pi\sigma_m^2}}{\sqrt{2\pi\sigma_e^2}\sqrt{2\pi\sigma_g^2}} (p) \exp\Big\{ -\frac{1}{2} \frac{\boldsymbol{r}_m^T \boldsymbol{r}_m}{\sigma_e^2} + \frac{1}{2\sigma_m^2} \mu_m^2 \Big\}}{\frac{\sqrt{2\pi\sigma_m^2}}{\sqrt{2\pi\sigma_e^2}\sqrt{2\pi\sigma_g^2}} (p) \exp\Big\{ -\frac{1}{2} \frac{\boldsymbol{r}_m^T \boldsymbol{r}_m}{\sigma_e^2} + \frac{1}{2\sigma_m^2} \mu_m^2 \Big\} + \frac{1}{\sqrt{2\pi\sigma_e^2}} (1-p) \exp\Big\{ -\frac{1}{2\sigma_e^2} \boldsymbol{r}_m^T \boldsymbol{r}_m^T \Big\}} \\
&\text{[break up terms over exp]} \\
&= \frac{\frac{\sqrt{2\pi\sigma_m^2}}{\sqrt{2\pi\sigma_e^2}\sqrt{2\pi\sigma_g^2}} (p) \exp\Big\{ -\frac{1}{2} \frac{\boldsymbol{r}_m^T \boldsymbol{r}_m}{\sigma_e^2} \Big\} \exp\{ \frac{1}{2\sigma_m^2} \mu_m^2 \Big\}}{\frac{\sqrt{2\pi\sigma_m^2}}{\sqrt{2\pi\sigma_e^2}\sqrt{2\pi\sigma_g^2}} (p) \exp\Big\{ -\frac{1}{2} \frac{\boldsymbol{r}_m^T \boldsymbol{r}_m}{\sigma_e^2} \Big\} \exp\{ \frac{1}{2\sigma_m^2} \mu_m^2 \Big\} + \frac{1}{\sqrt{2\pi\sigma_e^2}} (1-p) \exp\Big\{ -\frac{1}{2\sigma_e^2} \boldsymbol{r}_m^T \boldsymbol{r}_m^T \Big\}} \\
&\text{[common exp terms and constants from top/bottom]} \\
&= \frac{(p)\sqrt{\frac{\sigma_m^2}{\sigma_g^2}} \exp\Big\{ \frac{1}{2\sigma_m^2} \mu_m^2 \Big\}}{(p)\sqrt{\frac{\sigma_m^2}{\sigma_g^2}} \exp\Big\{ \frac{1}{2\sigma_m^2} \mu_m^2 \Big\} + (1-p)}
\end{aligned}
$$

In summary, to jointly sample $(\gamma_m, c_m)$, one first samples $c_m$. Then depending on if $c_m = 1$ we sample $\gamma_m$, and if $c_m = 0$ we set $\gamma_m$ to a point mass at 0:

$$
c_m \sim \text{Bern}(d_m)
$$

$$
\gamma_m \sim \begin{cases} \mathcal{N}(\mu_m, \sigma_m^2) & \text{if } c_m = 1 \\ 0 & \text{if } c_m = 0 \end{cases}
$$
